# Cerebral Blood Flow Increases Across Early Childhood

**DOI:** 10.1101/587139

**Authors:** Dmitrii Paniukov, R. Marc Lebel, Gerald Giesbrecht, Catherine Lebel

**Author notes:** Corresponding author: Catherine Lebel Room B4-513, Alberta Children’s Hospital 28 Oki Drive NW, Calgary, Alberta, Canada, T3B6A8.

## Abstract

Adequate cerebral blood flow (CBF) is essential to proper brain development and function. Detailed characterization of CBF developmental trajectories will lead to better understanding of the development of cognitive, motor, and sensory functions, as well as behaviour in children. Previous studies have shown CBF increases during infancy and decreases during adolescence; however, the trajectories during childhood, and in particular the timing of peak CBF, remain unclear. Here, we used arterial spin labeling to map age-related changes of CBF across a large longitudinal sample that included 279 scans on 96 participants (46 girls and 50 boys) aged 2-7 years. CBF maps were analyzed using hierarchical linear regression for every voxel inside the grey matter mask, controlling for multiple comparisons. The results revealed a significant positive linear association between CBF and age in distributed brain regions including prefrontal, temporal, parietal, and occipital cortex, and in the cerebellum. There were no differences in developmental trajectories between males and females. Our findings suggest that CBF continues to increase at least until the age of 7 years, likely supporting the ongoing improvements in behaviour, cognition, motor, and sensory functions of the children in early childhood.

**Highlights:**

* We mapped cerebral blood flow development longitudinally in 96 typically developing children aged 2-7 years
* Cerebral blood flow increased over the full age range
* No significant differences between males and females were found

**Conflicting of Interest:** R.M.L. is an employee of GE.

## 1. Introduction

Early childhood (2-7 years old) is an important period of development that sees large gains in cognitive, motor, and sensory function, as well as improvements in behaviour, all of which lay the foundation for future life. Changes in cognition and behaviour are supported by ongoing development of brain structure and function. During early childhood, the brain manifests shifts in functional connectivity (Long et al., 2017), cortical surface expansion (Remer et al., 2017), and increases in white matter volume (Deoni et al., 2012). Cerebral blood flow (CBF) may be particularly relevant to these changes, as it supplies blood and nutrients to the brain and supports ongoing development (Kandel et al., 2000). A non-invasive magnetic resonance imaging technique, arterial spin labelling (ASL), can measure CBF in the brain.

CBF is a measure of blood flow and perfusion, which underlies typical neuronal activity (Fox & Raichle, 1986) and cognitive function (Gur, et al., 1982). It also tends to be altered in disorders or diseases like schizophrenia (Pinkham et al., 2011), depression (Bench et al., 1992), anxiety (Gur et al., 1987), addiction (Wang et al., 2007), and head trauma (Wintermark et al., 2004b), and therefore may be a marker of abnormal development. Thus, understanding CBF changes in healthy children is critical for understanding both typical and atypical brain development.

Previous studies show that cortical CBF increases from birth to early childhood (Taki et al., 2011a) and decreases during adolescence (Avants et al., 2015; Satterthwaite et al., 2014). This suggests that there is a peak during early childhood, but the limited number of previous studies in this age range have produced conflicting results, showing ages of peak CBF that vary from 2 to 10 years (Carsin-Vu et al., 2018; Chiron et al., 1992; Liu et al., 2018; Takahashi et al., 1999; Wintermark et al., 2004a; Wu et al., 2016). Differences between studies are likely due, at least in part, to the very small number of participants included between ages 2-7 years (n=7-29) and the lack of longitudinal data, making it difficult to determine the actual trajectories during early childhood. Furthermore, calculating peak ages can be influenced by the inclusion of much older or much younger subjects, erroneously skewing the peak (Fjell, 2010). Therefore, a large longitudinal sample needs to be used to map the developmental trajectories of CBF.

Additionally, findings regarding sex differences have been very mixed. Most studies do not explicitly test sex differences. Among those that do, one study reported higher brain perfusion in parietal cortex in females than in males in participants 5-18 years old (Taki et al., 2011b), while another reported CBF decreases in males across adolescence, but a u-shaped trajectory of decreases then increases in female adolescents (Satterthwaite et al., 2014).

The goal of the current study was to characterize the developmental changes of CBF in a large, longitudinal sample of young children and evaluate sex differences.

## 2. Methods

### 2.1 Participants

Children were recruited from the Calgary area to participate in the study. Children underwent an MRI scan between the ages of 2-4 years and were invited to return for follow up imaging sessions approximately every 6 months. 105 children were initially recruited, providing 333 datasets total. One participant was excluded because of an incidental finding. All other datasets underwent a quality check by visual inspection, and datasets with excessive motion and/or scanning artifacts were excluded from the analysis, leaving 279 datasets on 96 children. These 96 children were 46 girls and 50 boys ranging in age from 1.97 to 6.90 years (M=4.46, SD=1.00 years); 80 were right-handed, 10 left-handed, and 6 with undetermined handedness. 32 children completed a single scan, 16 — two scans, 13 — three scans, 13 — four scans, 13 — five scans, 7 — six scans, 1 — seven scans, and 1 — ten scans. Parents provided written informed consent, and children provided verbal assent. The study was approved by the conjoint health research ethics board at the University of Calgary (REB18-1011).

### 2.2 Data acquisition

Neuroimaging data were collected using a research-dedicated GE 3T MR750w scanner (General Electric, Waukesha, WI) with a 32-channel head coil at Alberta Children’s Hospital. During the scan, children watched a movie of their choice. T1-weighted images were acquired using an FSPGR BRAVO sequence with TR = 8.23 ms, TE = 3.76 ms, inversion time of 540 ms, flip angle=12 degrees, voxel size = 0.9×0.9×0.9 mm^3^, 210 slices, matrix size=512×512, TI=540 ms, field of view=23.0. ASL images were acquired with the vendor supplied pseudo continuous 3D ASL sequence with TR = 4.56s, TE = 10.7ms, in-plane resolution of 3.5 mm, interpolated resolution of 1.73×1.73×4 mm^3^, post label delay of 1.5 seconds, and thirty 4 mm thick slices. The sequence scan time was 4.4 minutes.

### 2.3 Data preprocessing

ASL perfusion DICOM images were converted to CBF DICOM images on the scanner computer using the vendor supplied approach (consistent with Alsop et al., 2015). CBF is reported here in mL/100g/min.

T1-weighted and CBF images were converted from DICOM images to NifTi files using dcm2nii from the Mricron software package (Rorden & Brett, 2000). T1 images were registered to an asymmetric MNI template for 4.5-8.5-year-olds (Fonov et al., 2009, 2011) using N4-bias correction and nonlinear ANTs registration (Avants et al., 2009). The MNI template brain mask was transformed back to native T1 space for each subject, and T1 images were skull-stripped by multiplying the mask with the T1-image using FSL utilities (Jenkinson et al., 2012). The total volume of the brain was measured at this point. Each individual’s CBF image was registered to their T1 image using rigid transformation with 6 degrees of freedom as an estimator for the affine transformation, and with 6 degrees of freedom and the BBR cost function implemented in FSL’s FLIRT software (Greve and Fischl, 2009; Jenkinson & Smith, 2001, Jenkinson et al., 2002). Then, CBF images were transformed to MNI space using the transformation matrices from the steps above and skull-stripped using the MNI brain mask. CBF files were resampled to 2×2×2 mm using ANTs’ ResampleImage and concatenated into a single 4D file for further analysis.

### 2.4 Statistical Analysis

Statistical analysis was performed with hierarchical linear regression using Statsmodels (Seabold & Perktold, 2010) on every voxel inside the gray matter whole brain mask, which was a part of the pediatric MNI template (Fonov et al., 2009, 2011). To examine the significance of the estimated effects, we estimated cluster size threshold with 10 000 permutations using AFNI’s 3dClustSim (Forman et al., 1995) with non-Gaussian filtering, voxel-wise *p*-threshold of 0.01 for alpha of 0.05. The *t*-maps were thresholded at *t*=1.97 and merged to create final outputs with 3dmerge (Cox, 1996). All image preprocessing and analyses were scripted in Python 3.6 (Python Software Foundation, 2017) and bash (Free Software Foundation, 2007) utilizing Nipype software (Gorgolewski et al., 2011).

We analyzed CBF as a function of children’s age (including linear, quadratic, and cubic terms), sex, handedness, brain volume, and the interaction of age with sex. Brain volume was included to account for the head/brain growth that happens in children over this age range. To examine possible multicollinearity between age and brain volume, we computed the variance inflation factor, which equaled 1.09. Therefore, we concluded there was an absence of severe multicollinearity between the age and the brain volume.

### 2.5 Data and code availability

The preprocessing scripts along with analysis scripts and results are available online (https://github.com/dpaniukov/CBF_early_childhood). Research data are available upon request.

## 3. Results

### 3.1 Age-related changes in CBF

The quadratic and cubic terms for the children’s age were not significant, and therefore, were removed from further analyses. The final statistical model estimated the relationship between CBF and age, sex, age-by-sex interaction, brain volume, and handedness.

The results revealed a significant positive association between CBF and age, such that CBF increased as children got older, in prefrontal, temporal, parietal, and occipital cortex, and the cerebellum (see Figures 1-3). 11.5% of voxels in the grey matter mask showed significant increase of CBF with age, distributed across areas of the left frontal pole, left medial prefrontal cortex, left dorsolateral prefrontal cortex, bilateral superior frontal gyrus, left inferior frontal gyrus, bilateral premotor and motor regions, left parietal regions (supramarginal gyrus), left temporal regions (inferior, superior, and middle temporal gyri), bilateral occipital regions (lateral occipital cortex and occipital pole), and bilateral cerebellum. Across voxels with significant increases, the average estimated increase of CBF was 4.73 mL/100g/min per year, or 6.5% from baseline values across the age range. Across the whole gray matter mask, global CBF increased at a rate of 0.37 ml/100g/min per year, or 5.4% across the entire age range.

**Figure 1.**
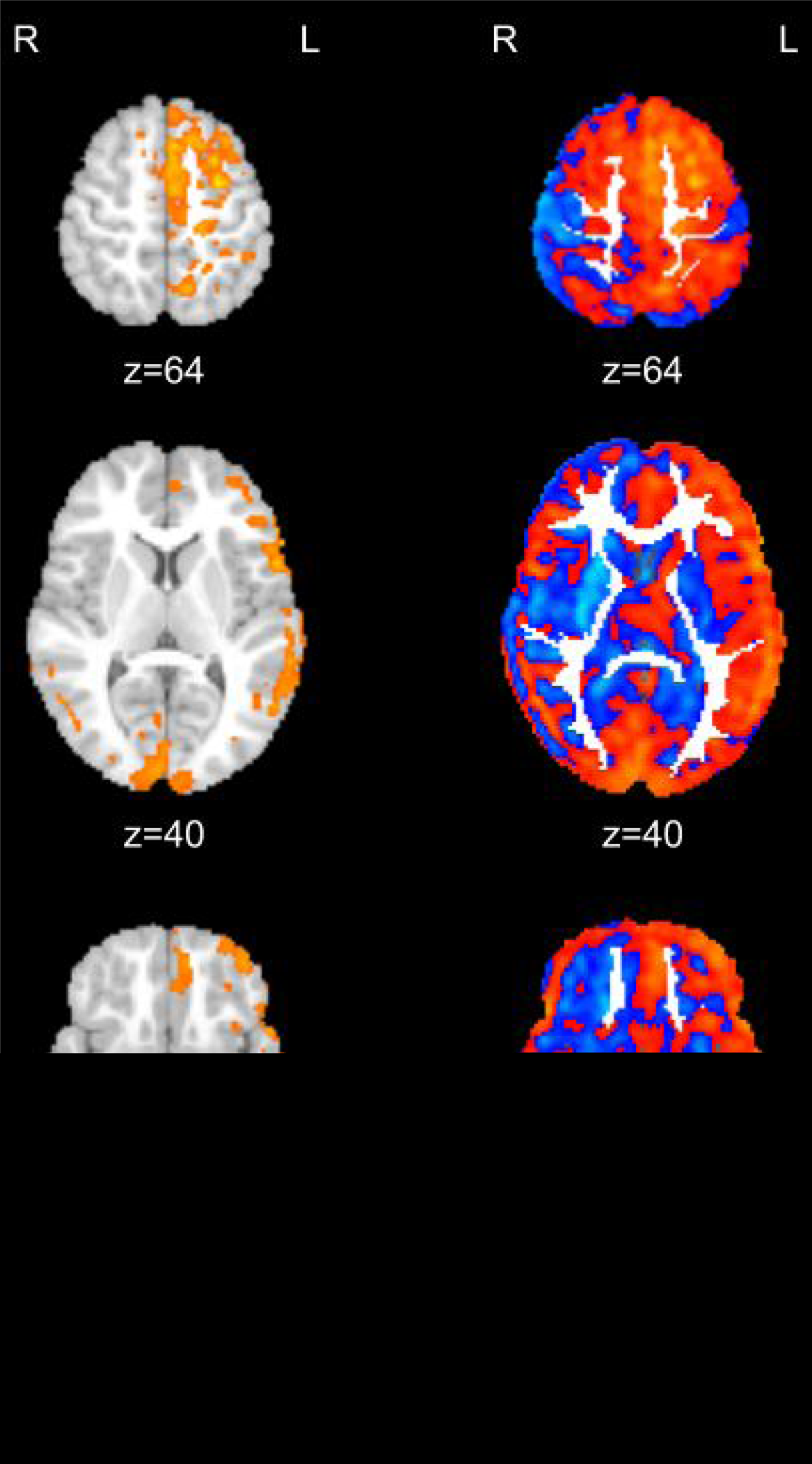
Relationship of CBF with age. The left column of images shows brain areas that had significant associations of CBF with age. The right column shows unthresholded t-statistics for the association of CBF with age. Red-yellow colors are for the increased association, blue-light blue colors are for the decreased association. The CBF increases globally in most of the regions of both hemispheres.

**Figure 2.**
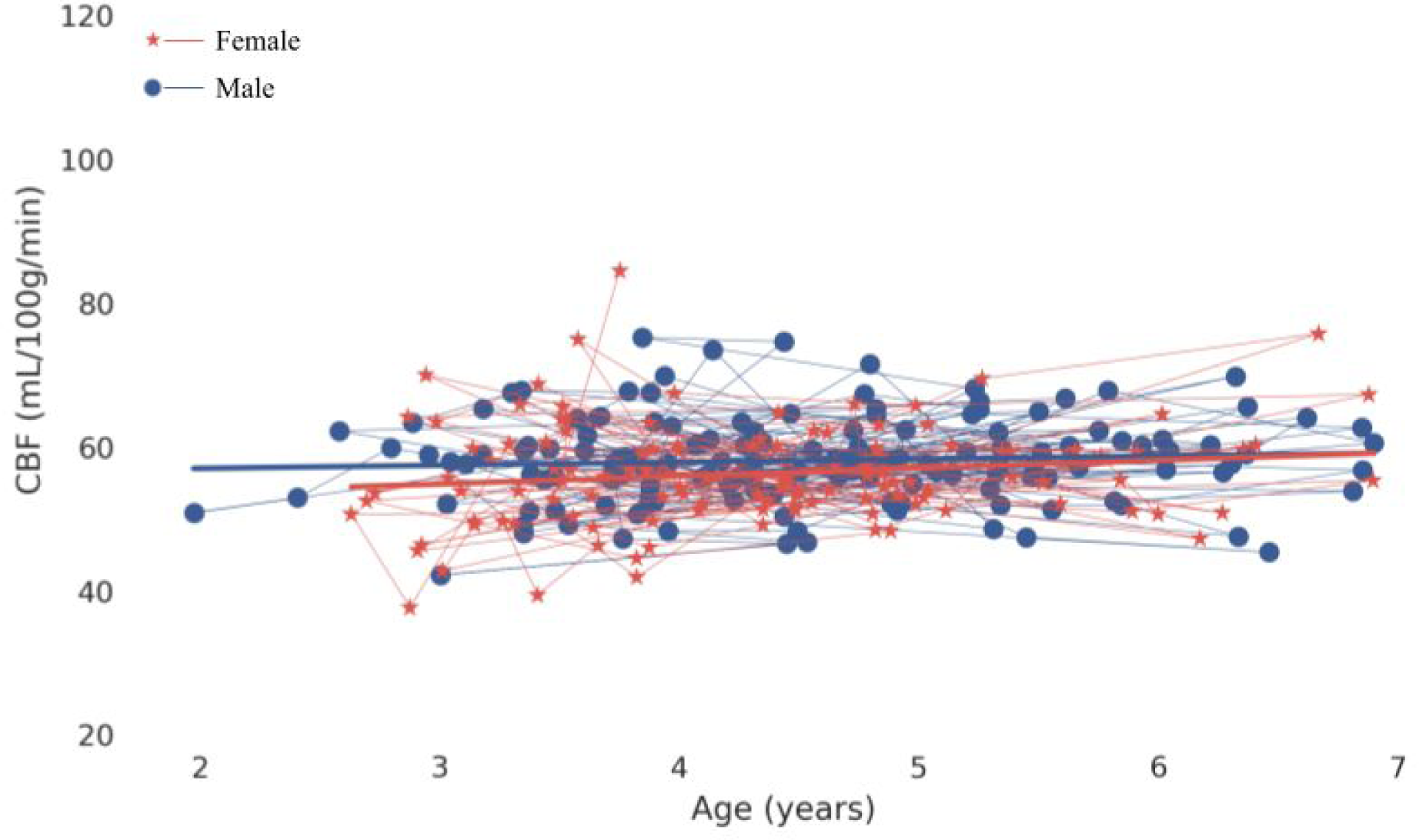
Global CBF increases with age. Each dot represents a scan, connected lines represent longitudinal scans, and the solid lines represent fits for males (blue) and females (red). Global CBF increased by 5.4% across the age range, 2-7 years.

**Figure 3.**
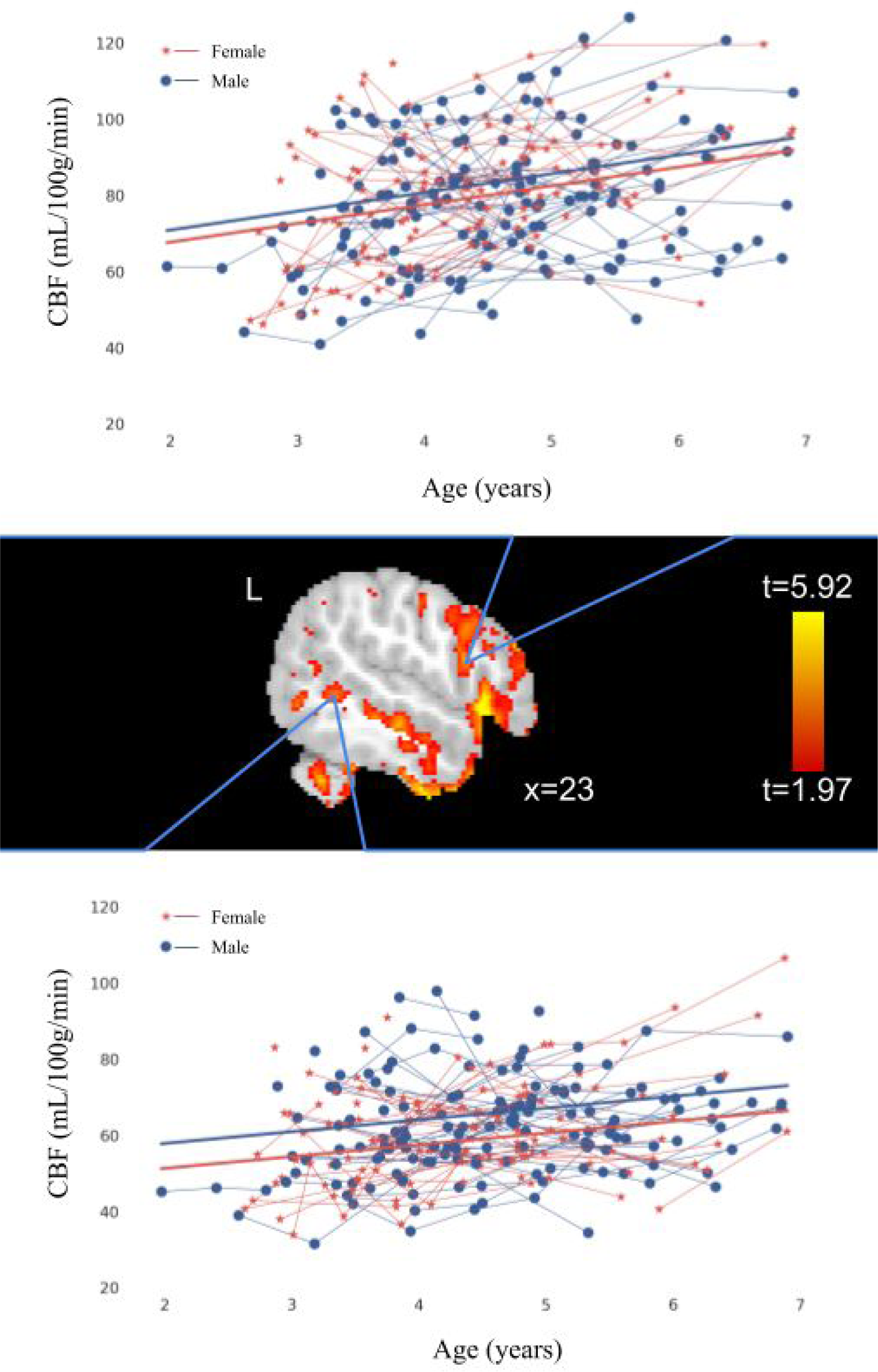
Relationship of CBF with age. These scatterplots are examples of areas with faster (top) and slower (bottom) rates of CBF change with age. The graphs show the increase of CBF with age for males (blue) and females (red) (no significant sex difference was found). Each dot represents a single scan time point; scans for the same participant are connected with lines that represent participants’ estimated trajectories.

### 3.2 CBF and sex differences

Females showed slightly lower CBF throughout the brain, but there were no significant associations of CBF with sex or with the interaction of age with sex.

## 4. Discussion

Here, in a large, longitudinal sample of young children, we show that CBF increases with age across the cortex and cerebellum in early childhood. Linear increases were seen from 2-7 years, with a mean increase of 6.5% over baseline values, or approximately 4.73 mL/100g/min per year. Trajectories of development were similar between males and females, and show that CBF continues to increase during childhood, suggesting peak values are obtained sometime after age 7 years.

Previous research found that CBF increases in infancy (Taki et al., 2011a), and almost doubles by 6 months of age, with less rapid increases continuing through infancy (Kehrer & Schöning, 2009); CBF is thought to peak between 2 and 10 years (Carsin-Vu et al., 2018; Chiron et al., 1992; Liu et al., 2018; Takahashi et al., 1999; Wintermark et al., 2004a; Wu et al., 2016), and decreases in adolescence (Avants et al., 2015; Satterthwaite et al., 2014). Peak ages have varied considerably across studies, likely because of small sample sizes and cross-sectional designs. Our results, showing slow but continuous linear increases of global CBF from 2-7 years, may indicate very active changes in connection of neurons that support ongoing development in cognition, sensory and motor functions, as well as behaviour.

CBF showed age-related increases across many brain areas. These areas are associated with semantic knowledge selection and attentional control (occipital, temporal, and parietal cortices; Thompson-Schill et al., 1997; Aron et al., 2003), language development and processing (left inferior frontal gyrus; Gabrieli et al., 1998) as well as with the evaluation of abstract information (rostrolateral prefrontal cortex; Vendetti & Bunge, 2014; Paniukov & Davis, 2018), and decision making (dorsolateral prefrontal cortex; Heekeren et al., 2006). These increases were distributed across the brain, all regions appearing to develop relatively similarly. The unthresholded t-map (Figure 1) shows some areas with negative associations with age, though these were all non-significant. That may imply that these regions are slowing down and perhaps nearing their peak CBF, though non-linear models were also nonsignificant for these regions, suggesting that peak CBF had not yet been achieved. Thus, the increases observed here appear to be more global in nature than concentrated in specific areas, but future research in other samples and/or with wider age ranges will be useful to pinpoint CBF trajectories in the areas not identified as significant here.

These changes in CBF occur in parallel with changes in brain structure and function During early childhood, brain structure also undergoes substantial development, including cortical thinning across cortex, cortical surface expansion (Remer et al., 2017), increases in white matter volume (Deoni et al., 2012), increases in fractional anisotropy, and decreases in mean diffusivity (Reynolds et al., submitted). Early childhood also sees substantial alterations in network connectivity (Long et al., 2017), when the brain reorganizes itself to support ongoing cognitive development. Increasing CBF may provide nutrients for the neurons in order to enable these structural and functional changes, and thus the significant behavioural and cognitive development during early childhood.

We did not find any significant sex differences in CBF or its developmental trajectories. One previous study in older children and adolescents (5-18 years) found higher brain perfusion in females than in males in the medial aspect of the parietal cortex (Taki et al., 2011b). Other studies showed more pronounced sex differences in CBF in adolescents with increase CBF in females (Satterthwaite et al., 2014). Interestingly, in an overlapping sample of children, we previously found similar fMRI measures of brain function between boys and girls (Long et al., 2017). Studies of cortical structure also suggest that sex differences may become more prominent later in development, with grey matter volume peaking earlier in males than females (Lenroot et al., 2007). Thus, it may be that CBF sex differences become more pronounced later in childhood or in adolescence, but are not apparent in early childhood.

This study has several limitations. To make scans comfortable and engaging for the children and to reduce potential head motion, children watched a movie of their choice during their scan. While patterns of functional connectivity from fMRI are broadly similar across passive viewing and “rest” conditions, it remains unclear whether this is true for CBF (e.g., Lim et al., 2010). That said, it is unlikely that our CBF values, which represent the average value over the 4.4 min acquisition, are strongly biased by the particular movie that a child chose to watch.

Therefore, future studies using rest conditions are also necessary to confirm associations between age and CBF, and studies measuring CBF during specific tasks may help further elucidate the role of CBF in cognitive development. Almost a third of the children did not provide longitudinal data. This relatively high attrition rate may have decreased the statistical power to detect effects, although the total sample size was still quite large.

## 5. Conclusion

For the first time using a large longitudinal sample, we show that CBF continues increasing until at least 7 years of age in distributed cortical and cerebellar regions. This suggests that peak CBF occurs sometime in later childhood, before the decreases previously observed in adolescence begin. Furthermore, we saw no differences in developmental trajectories between males and females, suggesting that such differences may arise later in childhood and adulthood. These increases of CBF likely support the ongoing development of cognitive, sensory, and motor functions.

## 6. Acknowledgements

This research was funded by CIHR (funding reference numbers IHD-134090, MOP-136797, MOP- 142387, and a New Investigator Award to C.L.), and an Owerko Postdoctoral Fellowship given to D.P. We would like to thank the APrON team at the University of Calgary for recruitment assistance.

